# HIF-stabilizing Biomaterials: from Hypoxia-mimicking to Hypoxia-inducing

**DOI:** 10.1101/2023.05.04.539396

**Authors:** Thibault Colombani, Khushbu Bhatt, Boris Epel, Mrignayani Kotecha, Sidi A. Bencherif

**Affiliations:** Department of Chemical Engineering, Northeastern University, Boston, MA 02115, USA; Department of Pharmaceutical Sciences, Northeastern University, Boston, MA 02115, USA; Department of Radiation and Cellular Oncology, The University of Chicago, Chicago, IL 60637, USA; Oxygen Measurement Core, O2M Technologies, LLC, Chicago, IL 60612, USA; Department of Bioengineering, Northeastern University, Boston, MA 02115, USA; Harvard John A. Paulson School of Engineering and Applied Sciences, Harvard University, Cambridge, MA 02138, USA; Biomechanics and Bioengineering (BMBI), UTC CNRS UMR 7338, University of Technology of Compiègne, Sorbonne University, 60203 Compiègne, France

**Keywords:** biomaterial, cryogel, cell culture, solid tumor, hypoxia, cancer

## Abstract

Recent advances in our understanding of hypoxia and hypoxia-mediated mechanisms shed light on the critical implications of the hypoxic stress on cellular behavior. However, tools emulating hypoxic conditions (i.e., low oxygen tensions) for research are limited and often suffer from major shortcomings, such as lack of reliability and off-target effects, and they usually fail to recapitulate the complexity of the tissue microenvironment. Fortunately, the field of biomaterials is constantly evolving and has a central role to play in the development of new technologies for conducting hypoxia-related research in several aspects of biomedical research, including tissue engineering, cancer modeling, and modern drug screening. In this perspective, we provide an overview of several strategies that have been investigated in the design and implementation of biomaterials for simulating or inducing hypoxic conditions—a prerequisite in the stabilization of hypoxia-inducible factor (HIF), a master regulator of the cellular responses to low oxygen. To this end, we discuss various advanced biomaterials, from those that integrate hypoxia-mimetic agents to artificially induce hypoxia-like responses, to those that deplete oxygen and consequently create either transient (< 1 day) or sustained (> 1 day) hypoxic conditions. We also aim to highlight the advantages and limitations of these emerging biomaterials for biomedical applications, with an emphasis on cancer research.

## 1. Biomaterial-mediated control of oxygen (O_2_) tension in cell culture

In biomedical research, tissue culture (or cell culture) has been instrumental in understanding cell biology and has led the way for numerous technological advancements and medical breakthroughs. The ability to grow cells in conditions like those found in their native environment provides the end user with a central platform for investigating several biological mechanisms, such as cellular processes, disease development, and drug responses.^1^ Cell culture also enables researchers to produce and engineer specific cell types relevant to various biomedical applications, including toxicity studies and cell-based therapies. Additionally, in the development of new treatments, it has been a valuable tool for investigating the effects of environmental factors on cells, such as genetic mutations.^2^

With the recent findings on the impact of low O_2_ tension on cellular behavior, emulating hypoxic conditions in cell culture has become a critical aspect of *in vitro* research.^3,4^ For instance, hypoxia, which is a low level of O_2_ (< 3–5%) in tissues, is a common feature of solid tumors and a hallmark of cancer.^4–6^ Specifically, solid tumors are abnormal masses or lumps formed by the growth of malignant cells in various parts of the human body (***Figure 1A***). They are usually heterogenous and characterized by normoxic, necrotic, and hypoxic regions (***Figure 1B***).^7^ The normoxic region comprises actively growing and functioning cells that continue to proliferate. The necrotic region typically found in the inner core of solid tumors is characterized by widespread cell death. The hypoxic region, defined by insufficient O_2_ supply due to vascular insufficiency, exerts a profound effect on tumor cells, resulting in metabolic changes, uncontrolled proliferation and survival, and increased angiogenesis.^4^ This environment has been associated with an increased risk of metastasis, cancer progression, and therapy resistance. Since tumor hypoxia is a predictor of poor clinical outcomes across all cancers, a better understanding of the effects of hypoxic stress on cancer cells, including phenotypic changes and tumor heterogeneity, would help develop more effective therapies.^8^ Therefore, by creating controlled hypoxic conditions that simulate the tumor microenvironment, the effectiveness of new drugs and treatments could be tested more efficiently and reliably.^9^ This strategy could eventually improve the accuracy of preclinical studies and ultimately reduce the high failure rates in late-phase clinical trials.^10^

**Figure 1.**
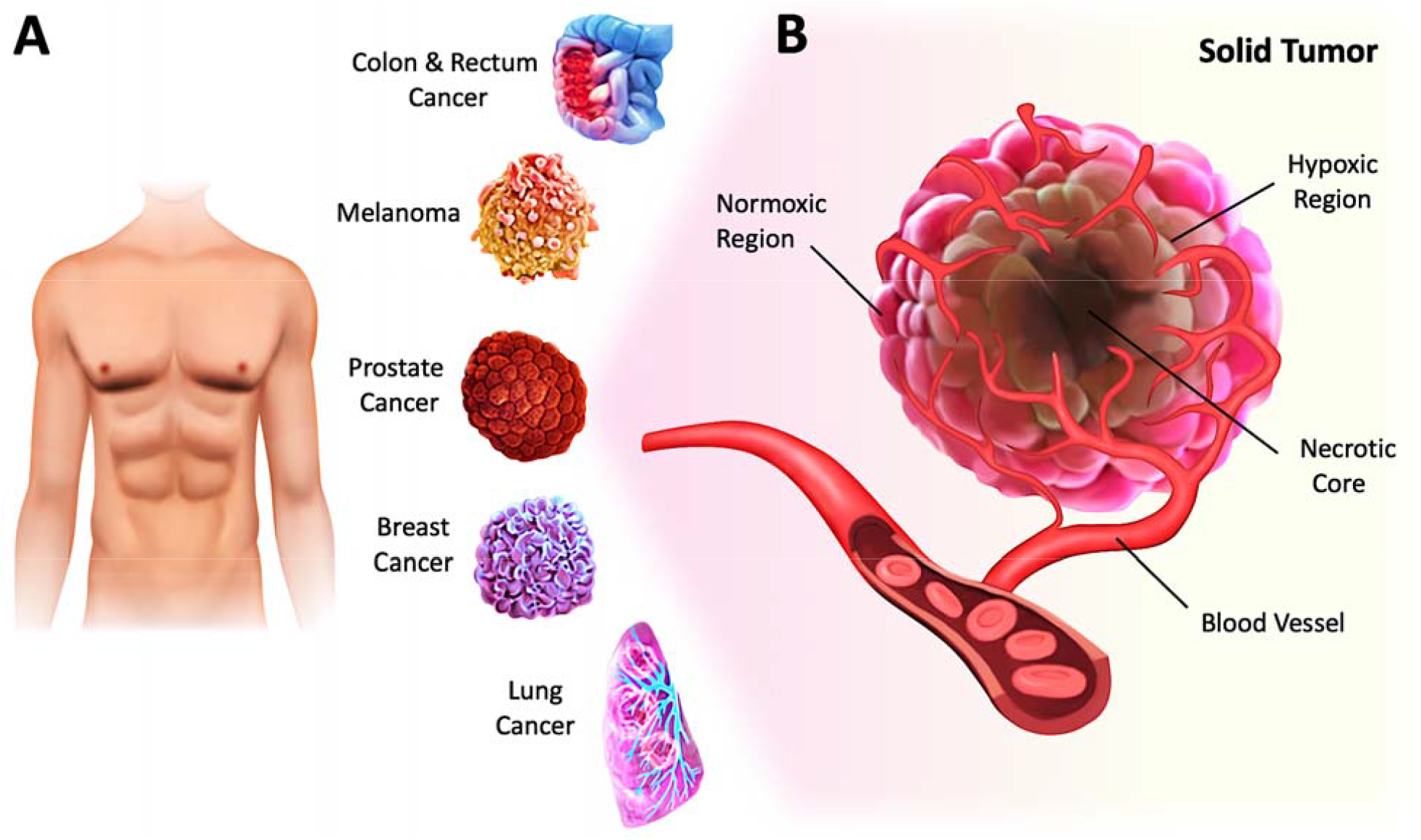
Hypoxia is a hallmark of cancer. **A**) Solid tumors account for approximately 90% of adult human cancers and can develop in many parts of the human body, including breast, lung, prostate, colon, rectum, and skin. **B**) Illustration of a solid tumor harboring profound chronic hypoxia and necrosis in regions distant from the vasculature.

Hypoxia-inducible factor-α (HIF-α), a transcription factor stabilized under hypoxic stress, plays a central role in the regulation of cellular responses.^11,12^ Upon exposure to hypoxia, the restrained enzymatic activity of PHD results in the rapid stabilization and accumulation of HIF-α (i.e., HIF-1α, HIF-2α, HIF-3α) in the cytoplasm. In this process, O_2_-dependent PHD-mediated hydroxylation, pVHL-dependent ubiquitination, and proteasomal degradation of HIF-α are all suppressed (***Figure 2***, *left panel*). As a result, accumulated HIF-α translocates to the nucleus, where it dimerizes with HIF-1β, and the activation complex subsequently binds to HRE sequences on DNA. The recruitment of the transcriptional co-activators p300/CBP upregulates hundreds of target genes involved in multiple signaling pathways, including cell survival and proliferation, angiogenesis, metabolic reprogramming, and drug resistance (***Figure 2***, *right panel*). Due to the nature of solid tumors to develop regions permanently or transiently subjected to hypoxia, a better understanding of the role of HIF-α in cancer development and progression is essential for advancing the field and discovering more effective therapies.

**Figure 2.**
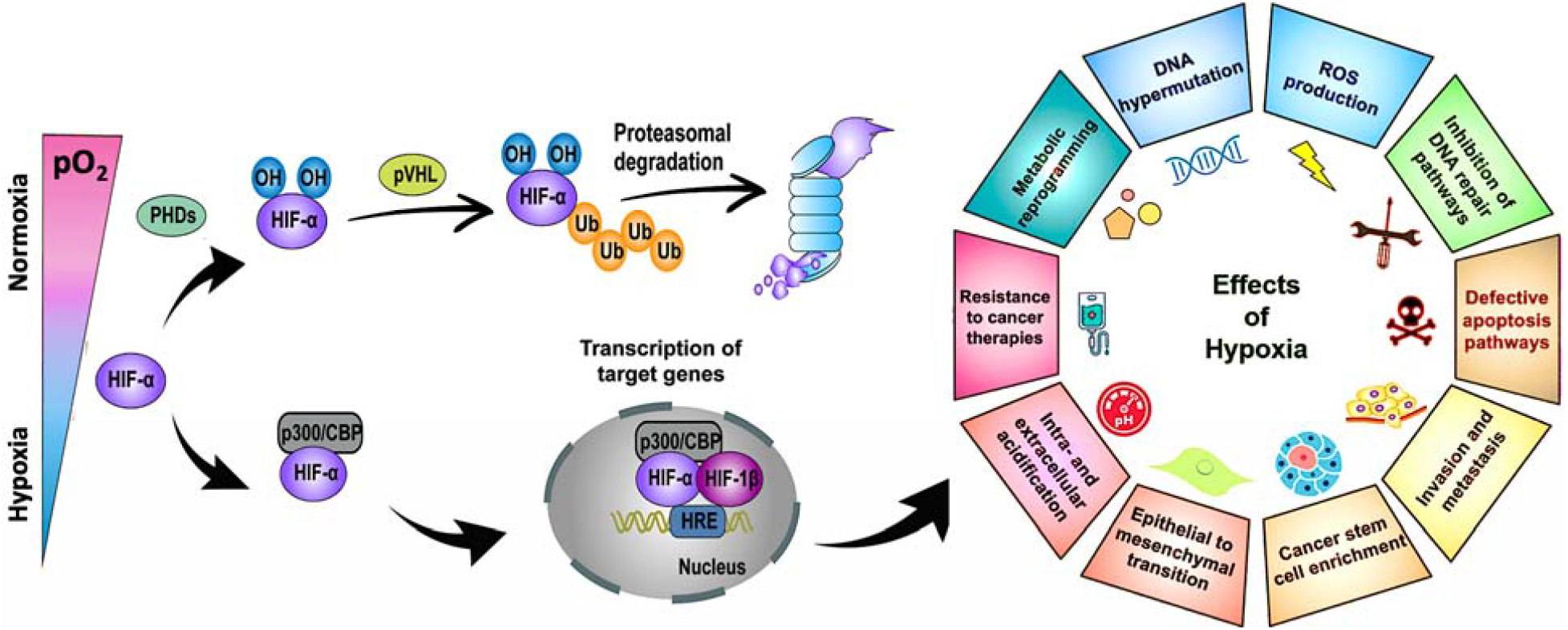
Stabilization of HIF-α at low O_2_ tension plays a crucial role in regulating the cellular responses of tumors to hypoxia. Schematic of the regulation of HIF-α (i.e., HIF-1α, HIF-2α, HIF-3α), its transcriptional activity, and the effects of hypoxia in cancer development and progression. PHD: prolyl hydroxylase. CBP: CREB (cyclic adenosine monophosphate response element binding protein) binding protein. Ub: ubiquitin. pVHL: Von Hippel–Lindau tumor suppressor. HRE: hypoxia-response element. ROS: reactive oxygen species. DNA: deoxyribonucleic acid. Reproduced with permissions from ref. 9 and 10. http://creativecommons.org/licenses/by/4.0/

As evidence increasingly indicates that O_2_ strongly and undeniably influences and regulates cellular behavior, we are witnessing a paradigm shift in the field of tissue engineering. More scientists and researchers are turning to *in vitro* cell culture under hypoxic conditions, enabling the translation of findings under pathological conditions *in vitro* to diseases. However, commonly used hypoxia-inducing methods (i.e., tri-gas incubators, chemical inhibitors, hypoxia-mimetic agents) for artificially triggering bona fide hypoxic responses in cell culture have shown a number of limitations.^13,14^ These techniques are often unreliable and/or often result in off-target effects, thereby inducing inconsistent and uncontrolled hypoxic conditions. Furthermore, these methods could be cytotoxic and interfere with normal cellular processes, pinpointing the need to investigate alternative strategies to induce hypoxia. Creating a more controlled hypoxic environment in cell culture can allow for accurate study of the effects of hypoxia on tumor cells and result in fewer disparities or inconsistencies between studies. These alternative strategies could also provide new avenues for studying hypoxia and its role in biological processes. To address this issue, many researchers are inclined to explore hypoxia-inducing biomaterials to create a more suitable cell culture environment. Hypoxia-inducing biomaterials are a class of materials used to mimic the low O_2_ tension environment found in many physiological and pathological processes. These biomaterials are designed to promote hypoxia *in vitro* and can provide a controlled and reproducible environment for studying cellular responses at low O_2_ levels.

Hydrogels, which were among the first materials ever developed for human use, have been extensively investigated. Due to their crosslinked polymeric network and high-water content, hydrogels are deemed safe, display excellent performance, and provide a physiologically relevant environment for cells.^15^ As they resemble soft tissues, hydrogels have been used as three-dimensional (3D) scaffolds that mimic the extracellular matrix (ECM) of native tissues. Furthermore, hydrogels can be easily functionalized with biomolecules such as peptides and proteins to support cell adhesion, proliferation, and differentiation. As the field of biomaterials is constantly evolving, recent advances in the design of sophisticated hydrogels with the ability to induce hypoxia have the potential to change the way we perform cell culture, advance our understanding of cell biology, and ultimately develop new therapeutic strategies, especially in cancer.^16^

In this perspective, we first provide some insights into the potential and utility of biomaterials to mimic or create hypoxic conditions for cell culture. Next, we dive into the various strategies that have been investigated, from integrating hypoxia-mimetic agents to fabricating hypoxia-inducing biomaterials. As a platform technology, such approaches can be leveraged to study and better understand many cellular processes (e.g., angiogenesis, cellular survival, tumorigenesis) involved in cancer initiation and development or to screen and identify anticancer therapeutics more reliably.

## 2. Hypoxia-mimetic agents to artificially trigger hypoxia-like responses

Hypoxia-mimetic agents are molecules that mimic the effects of hypoxia by stabilizing HIF-1α on a cellular level.^17^ This allows its translocation to the nucleus, wherein it heterodimerizes with HIF-1β to regulate the expression of over 100 genes involved in cell growth and differentiation, metabolism, apoptosis, erythropoiesis, and angiogenesis.^18^ The most commonly used strategy relies on inhibiting PHDs using iron chelators and competitors, 2-oxoglutarate (2-OG) analogs, or nitric oxide donors.^17^ Iron chelators such as deferoxamine mesylate (DFO) and iron competitors such as Co^2+^ in the form of cobalt chloride (CoCl_2_) are widely used to reduce the amount of free Fe^2+^ available for catalyzing hydroxylation of HIF. 2-OG analogs such as dimethyloxalylglycine (DMOG), hydroxybenzoic acids, and hydroxybenzenes prevent HIF degradation by inhibiting PHDs via competing with PHD substrates such as 2-OG.^19^ These agents overcome the need to deplete O_2_ while stabilizing HIF-1α, thereby promoting angiogenesis and tissue regeneration. Nevertheless, hypoxia-mimetic molecules can induce cytotoxicity over long-term exposure.^19^

To address these challenges, hypoxia-mimicking biomaterials have been designed to incorporate hypoxia-mimetic agents to simulate hypoxic conditions while also improving their release kinetics and cytocompatibility. For instance, DFO has been loaded in a biomaterial scaffold to prolong its duration of action and decrease its cytotoxicity.^20^ Additionally, the scaffold provided a hypoxic environment conducive to promoting bone and tissue regeneration, angiogenesis, and immunomodulation. In another study, 3D-nanofibrous gelatin scaffolds incorporating mesoporous silica nanoparticles (MSNs) have been designed to deliver bone morphogenetic protein (BMP-2) and DFO for inducing angiogenesis and osteogenesis.^21^ BMP-2 encapsulated in large-pored MSNs led to its sustained release, whereas covalent conjugation of chitosan to DFO promoted its short-term release and improved half-life, thereby reducing its cytotoxic potential. Similarly, hydrogel scaffolds, with a collagen-based core containing BMP-2 and an alginate-based shell encapsulating Co^2+^ ions, have been fabricated to allow rapid release of Co^2+^ and sustained delivery of BMP-2.^22^ Other approaches have investigated the use of PHD inhibitors to enhance the angiogenic properties of biomaterials that inherently possess osteogenic activities. For instance, several groups combined bioactive glass scaffolds with Co^2+^ or Cu^2+^ for bone tissue engineering and dental stem cell regeneration.^23–26^ In addition, multi-walled carbon nanotubes (MWCNTs) were combined with Co^2+^ nanocomposites to promote the sustained release of Co^2+^, thereby maintaining hypoxia-like conditions for a longer period. Since MWCNTs possess good electrical conductivity, they are promising candidates for cardiac, neural, and muscle tissue engineering.^27^

Multiple groups have investigated the encapsulation of mesenchymal stem cells (MSCs) in hydrogel scaffolds containing DMOG to achieve enhanced retention, controlled release, and targeted delivery for bone restoration.^28,29^ For instance, encapsulated DMOG and MSCs in alginate-based hydrogels functionalized with arginyl–glycyl–aspartic acid (RGD) allowed their short-term release to stabilize HIFs and promoted chondrogenesis of MSCs compared to DMOG-free alginate.^28^ Conversely, DMOG-loaded polycaprolactone grafts improved the sustained release of DMOG, facilitating the proliferation and migration of human umbilical vein endothelial cells in a rat abdominal artery model. In addition, they promoted local M2 macrophage polarization, the release of pro-angiogenic factors, and the contractile phenotype of smooth muscle cells, leading to enhanced vascular repair and modulation of graft-induced inflammation.^30^

Overall, the combination of hypoxia-mimetic agents and biomaterials has been shown to improve the release kinetics, extend the duration of action, induce the targeted delivery, and increase the cytocompatibility of these chemicals. However, the cytotoxicity of such molecules needs to be evaluated more thoroughly, as their effect is usually limited to the HIF-1α subunit, and they cannot realistically emulate the complexity of tissues with distinct levels of hypoxia.

## 3. Emerging biomaterials: O_2_-depleting hydrogels to induce hypoxia

O_2_-depleting biomaterials have been investigated to remediate the limits of chemically induced hypoxia while recapitulating the biochemical and biophysical complexity of the natural ECM. This led to the design of hypoxia-inducible hydrogels which rely on the consumption of O_2_ via the chemical crosslinking of the polymer networks.^31–33^ These hydrogels have been fabricated with naturally derived polymers to display biological properties (e.g., gelatin) and chemically modified with ferulic acid (FA). In the presence of laccase, FA molecules dimerize by oxidation, resulting in the crosslinking of FA-containing polymers and the depletion of O_2_. Hypoxia-inducible hydrogels have been shown to induce hypoxia within 10 min, which lasted for 12 h in a cell-free environment and up to 48 h with encapsulated endothelial-colony-forming cells with minimal toxicity.^31,33^ Such biomaterials can be used to study the impact of hypoxia and matrix viscoelasticity on the process of angiogenesis,^32^ or more broadly to investigate new biological pathways triggered in O_2_-deficient environments for regenerative medicine, disease modeling, or cancer studies. However, hypoxia-inducible hydrogels have several limitations. They typically induce transient hypoxia and, as a result, expose cells to reoxygenation, leading to short-lived stabilization of HIF-1α and HIF-2α (≤ 12 h).

Since FA-based systems only allow for short-term hypoxia investigations, efforts have been made to develop hypoxia-inducing biomaterials for long-lasting O_2_ depletion.^34–36^ This approach relies on enzymatic reactions that continuously deplete O_2_, thus creating more sustained hypoxic conditions for cell culture. The most notable enzyme is glucose oxidase (GOX) which oxidizes β-D-glucose, commonly present in cell culture media, into gluconic acid and hydrogen peroxide (H_2_O_2_) while consuming O_2_ in the process. Due to H_2_O_2_ cytotoxicity, catalase (CAT) is commonly added to the cell culture media. CAT is a potent enzyme that facilitates the breakdown of H_2_O_2_ into H_2_O and ½ O_2_. While the enzymatic GOX/CAT duo has been used for over a decade to induce hypoxia in aqueous solutions,^37^ its short-lived stability has resulted in transient hypoxia. To address this issue, chemically modified GOX has been first immobilized in hydrogels to fabricate hypoxia-inducing hydrogels (HIHs).^35^ C. Lin and colleagues have demonstrated that GOX-grafted poly(ethylene glycol) diacrylate (PEGDA-GOX) hydrogels were capable of maintaining hypoxic conditions within the cell culture media for up to 24 h while also retaining high cell viability and inducing increased expression of hypoxia-associated gene carbonic anhydrase 9 (Ca9). However, to reduce H_2_O_2_-associated cytotoxicity, this system still required the addition of soluble CAT into the media. Furthermore, cell reoxygenation occurred after 48 h, and only ∼55% of acute myeloid leukemia (Mol14) cells remained viable after 2 days. To extend hypoxia lifetime and decrease cytotoxicity, the same team has also investigated the addition of glutathione (GSH), a potent and biocompatible antioxidant, in their cell culture media to reduce H_2_O_2_.^36^ As a toxic by-product, H_2_O_2_ not only impacts cell viability but also inactivates the enzymatic activity of GOX/CAT. The presence of GSH further extended the hypoxic conditions induced by HIHs for up to 48 h. However, the cytotoxicity towards Mol14 cells remained a major concern. While these advances were disruptive, there is still a need to design more cutting-edge biomaterials that can immobilize both enzymes, enabling sustained, controlled, and long-lasting hypoxia, and that will ultimately be safer for mammalian cells and preserve high cell viability and function. In addition, such alternatives could circumvent the inherent shortcomings of standard hydrogels, including a mesoporous structure that results in hampered cell infiltration, poor diffusion of O_2_ and nutrients, ineffective removal of cellular waste, and inadequate biophysical properties.

## 4. Hypoxia-inducing cryogels (HICs) to induce long-standing hypoxia

Recently, a new class of injectable hydrogels known as cryogels has emerged as a promising biomaterial for tissue engineering, cell transplantation, tumor modeling, and immunotherapies.^38–41^ Cryogels possess unique features such as shape memory properties, high viscoelasticity, and large interconnected pores that address conventional hydrogels’ mass transport and diffusion limitations. Cryogels’ unique architecture allows tumor cell reorganization in 3D and immune cell infiltration, enabling the development of more relevant tumor models. In addition, our team has demonstrated their straightforward functionalization with peptides, proteins, and enzymes,^41,42^ making them an ideal platform for cancer and immunology research as well as for designing long-standing hypoxia-inducing biomaterials.

In our latest work, our team engineered hypoxia-inducing cryogels (HICs) to create 3D microtissues with features of the hypoxic tumor microenvironment (TME).^43^ Cryogels were fabricated at subzero temperature with chemically modified hyaluronic acid (HAGM), an essential component of the tumor-associated ECM, and RGD, a short peptide found in various proteins of the ECM. In addition, the scaffolds were grafted with immobilized GOX and CAT to deplete O_2_ rapidly and safely (***Figure 3***). HICs induced hypoxia within 1 h both inside the scaffolds and in their surrounding environment (***Figure 4***). Strikingly, they maintained hypoxia for at least 6 days while remaining cytocompatible and biocompatible,^43^ which is a significant improvement compared to HIHs. HICs also induced sustained cellular hypoxia and facilitated the reorganization of cancer cells (4T1 mammary carcinoma and B16-F10 melanoma) into microsized tumor masses.

**Figure 3:**
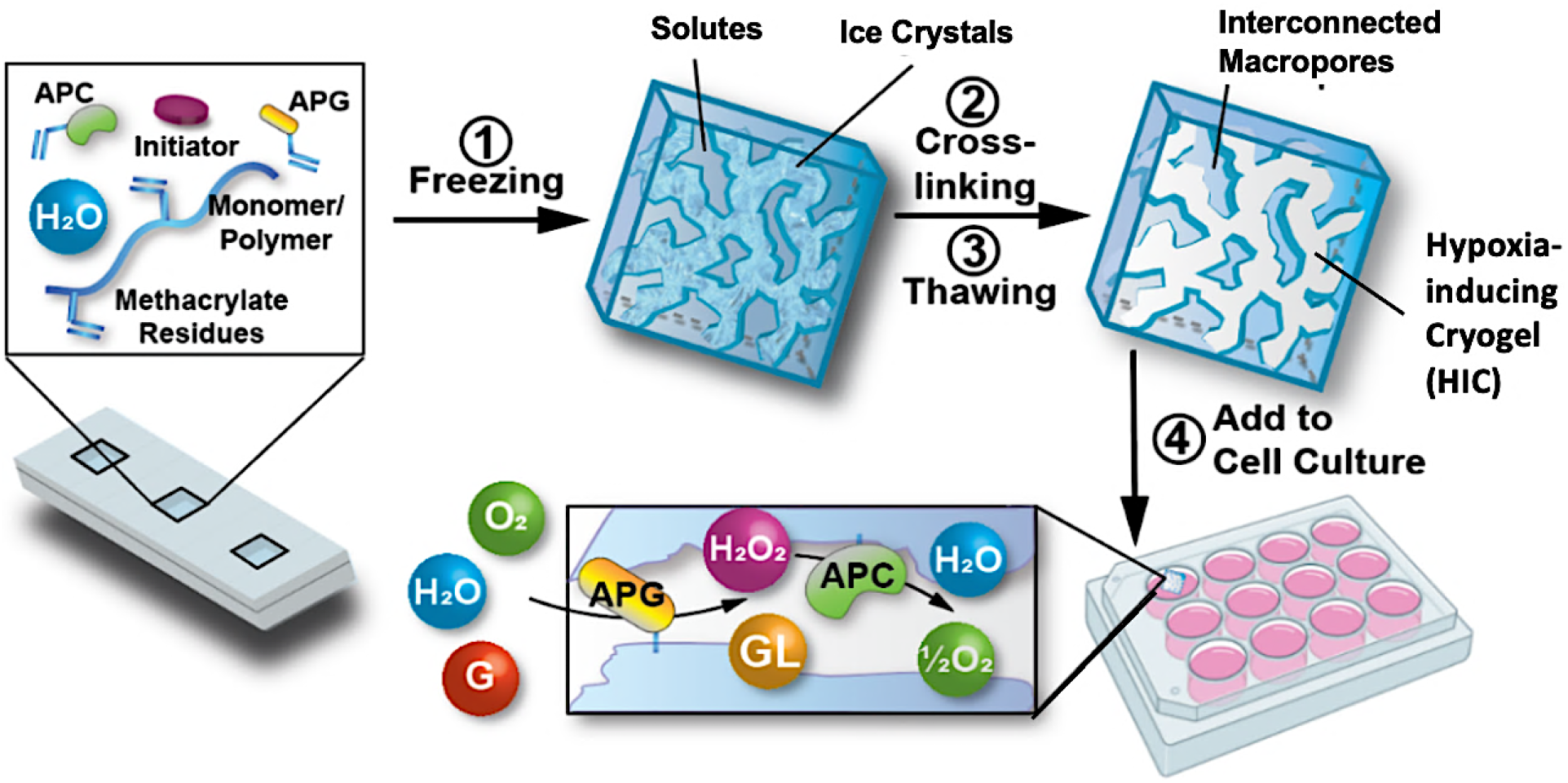
Schematic illustration of the process of fabricating HICs and their ability to induce a hypoxic microenvironment for cell culture. Hypoxia-inducing cryogels (HICs) are fabricated by mixing methacrylated hyaluronic acid (HAGM), acrylate-PEG-RGD, acrylate-PEG-glucose oxidase (APG), acrylate-PEG-catalase (APC), and the initiator system (TEMED/APS) in an aqueous solution and then allowing it to cryopolymerize at subzero temperature (−20°C). Once the cryogelation is complete, HICs are thawed at room temperature and start consuming O_2_ when immersed in a cell culture medium. APG depletes O_2_ by converting D-glucose (G) to hydrogen peroxide (H_2_O_2_) and D-glucono-δ-lactone (GL), while APC breaks down H_2_O_2_ into water (H_2_O) and ½ O_2_, inducing sustained hypoxic conditions. Reproduced with permission from ref. 43.

**Figure 4:**
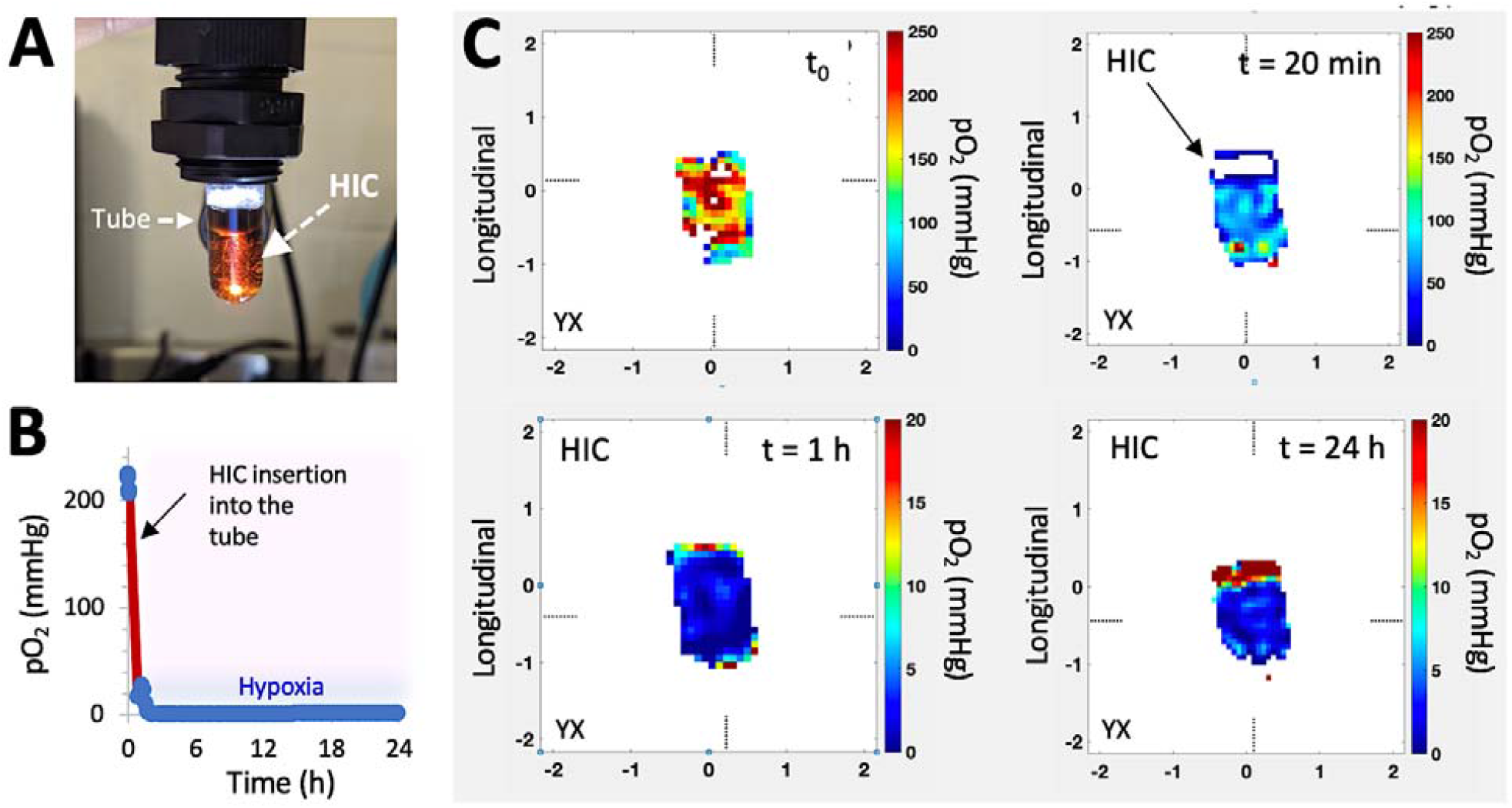
HICs deplete O_2_ quickly and induce sustained hypoxia. **A–C**) Characterization of O_2_ depletion and mapping of hypoxia using electron paramagnetic resonance oxygen imaging (EPROI). **A**) Digital image showing a HIC inserted in a glass tube for the EPROI test. O_2_ mapping was performed on a 25 mT EPR imager (JIVA-25™, O2M Technologies, LLC, Chicago, IL). A cylindrical HIC (6 mm diameter, 5 mm height) was inserted in a 10-mm diameter test tube and submerged with 1 mL of PBS supplemented with D-glucose (4.5 g/L) and trityl radical molecular probe (1 mM OX071) at 37 °C. **B**) Average pO_2_ measurements in the whole system (solution and HIC) *vs*. time. **C**) pO_2_ distributions on the tangential plane of a HIC at different time points.

HICs also demonstrated their ability to promote hypoxia-mediated cancer cell metabolism and regulation of gene expression. For instance, B16-F10 cells cultured in HICs switched their metabolism from aerobic respiration to anaerobic glycolysis, leading to decreased pH and glutamine concentration in cell supernatant which were correlated with increased secretion of glutamate and lactate. Similarly, they induced the upregulation of HIF-1α, VEGFα, and SOX2 genes associated with tumor vascularization and tumor cell proliferation, migration, and invasion. Finally, these cancer cells also displayed increased resistance to commonly used chemotherapeutics (doxorubicin and cisplatin) and enhanced sensitivity to tirapazamine, a hypoxia-activated drug. Altogether, this set of data demonstrated the potential of HICs as a platform to model hypoxic solid tumors and study hypoxia-mediated resistance to anticancer therapies (***Figure 5***).

**Figure 5:**
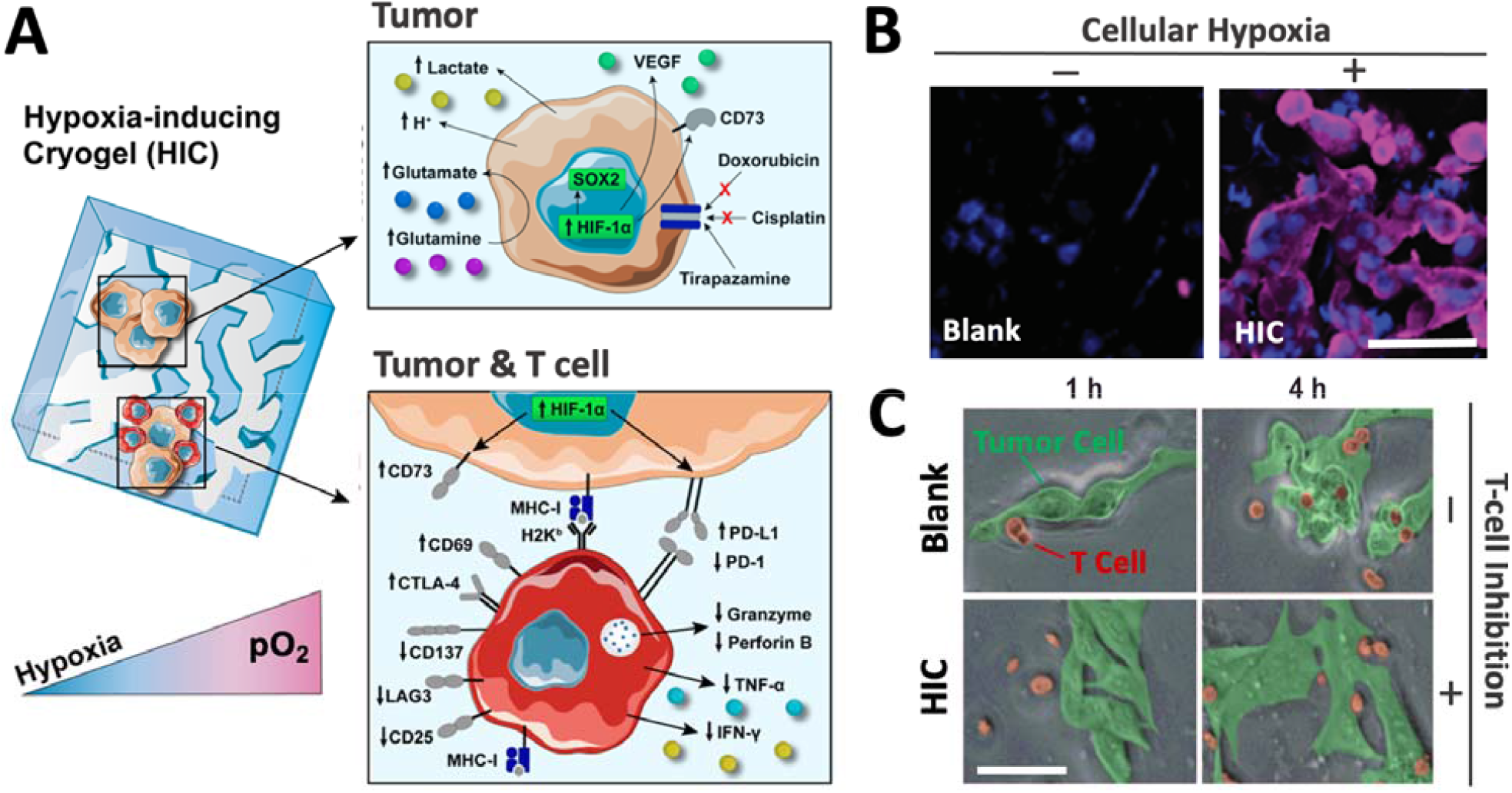
HICs recreate a hypoxic tumor microenvironment and can be leveraged as a tool to investigate the effects of hypoxia on tumors and immune cells. **A**) Adaptive changes of cellular metabolism, phenotype, and treatment response when tumor cells are cultured in HICs (upper panel). HICs can also be utilized to mimic tumor–immune cell interactions and unravel hypoxia-driven immunosuppression mechanisms on tumor-reactive T cells (lower panel). **B**) Confocal images depicting cellular hypoxia in B16-F10 cells cultured in blank cryogel and HIC for 24 h. Blue staining (DAPI) = nuclei; purple staining (Hypoxyprobe-1) = cellular hypoxia. **C**) Time-lapse confocal images depicting T-cell-mediated attack against B16-OVA cells in enzyme-free cryogels (blank) and HICs after 1 and 4 h of co-culture. Scale bars = 50 μm. Reproduced with permission from ref. 43.

Finally, our team has recently leveraged the HIC technology to better understand cancer immunology. We demonstrated that HICs allowed immune cell infiltration while recapitulating the hypoxia-mediated inhibition present in hypoxic TMEs. We confirmed hypoxia-mediated dendritic cell (DC) immunosuppression and highlighted the dual implication of both low O_2_ tension and ECM composition on DC phenotype and inhibition. In addition, we showed that HIC-based cancer immunology models could recapitulate the inhibition of cytotoxic T-cell (CTL) killing in hypoxic conditions. Specifically, we observed that CTLs co-cultured with cancer cell–laden HICs displayed a significant decrease in pro-inflammatory cytokine secretion (IFN-γ and TNF-α), an increased expression of the checkpoint inhibitor CTLA-4, and a lack of cytotoxic protein degranulation (perforin and granzyme) responsible for cytotoxic T-cell function (***Figure 5***). New platforms such as HICs could potentially improve the efficiency of anticancer drug screening and discovery and further expand our understanding of cancer immunology. By providing a closer inspection of hypoxic conditions, they may help unravel new mechanisms associated with tumor-associated immunosuppression and ultimately continue to push the boundaries of cancer immunotherapy.

## 5. Future perspective and conclusions

Despite its clinical importance, conducting hypoxic cell culture remains a major hurdle. Thankfully, state-of-the-art technologies are being developed at a rapid pace and should further change the way we perform tissue culture. As discussed in this perspective, biomaterials have a pivotal role to play in establishing prolonged or intermittent hypoxic conditions due to their versatility and popularity in the field. For instance, hypoxia-inducing gel systems with immobilized enzymes, and featuring ECM-like composition and microstructure, are set to advance the field of cancer research further and democratize hypoxic cell culture. Current and emerging technological advances should open new avenues for the design of sophisticated research tools for studying cells and tissues subjected to hypoxic stress. These advances should also deepen our understanding of the molecular mechanisms of hypoxia in the pathogenesis of various diseases, including cancer, myocardial ischemia, and endometriosis.

## 7. Acknowledgments

The authors thank the Institute for Chemical Imaging and Living Systems (ICILS) for their technical assistance with the confocal microscope. S.A.B. gratefully acknowledges the financial support from the National Institutes of Health (NIH, 1R01EB027705) and the National Science Foundation (NSF CAREER, DMR 1847843). Zachary Rogers at Northeastern is acknowledged for his contribution to sample preparation for the EPR O_2_ analysis and imaging.

## Table of contents image

Breakthroughs in biomaterials science open new avenues for the stabilization of hypoxia-inducible factor-α (HIF-α), a master regulator of the cellular responses to low oxygen.

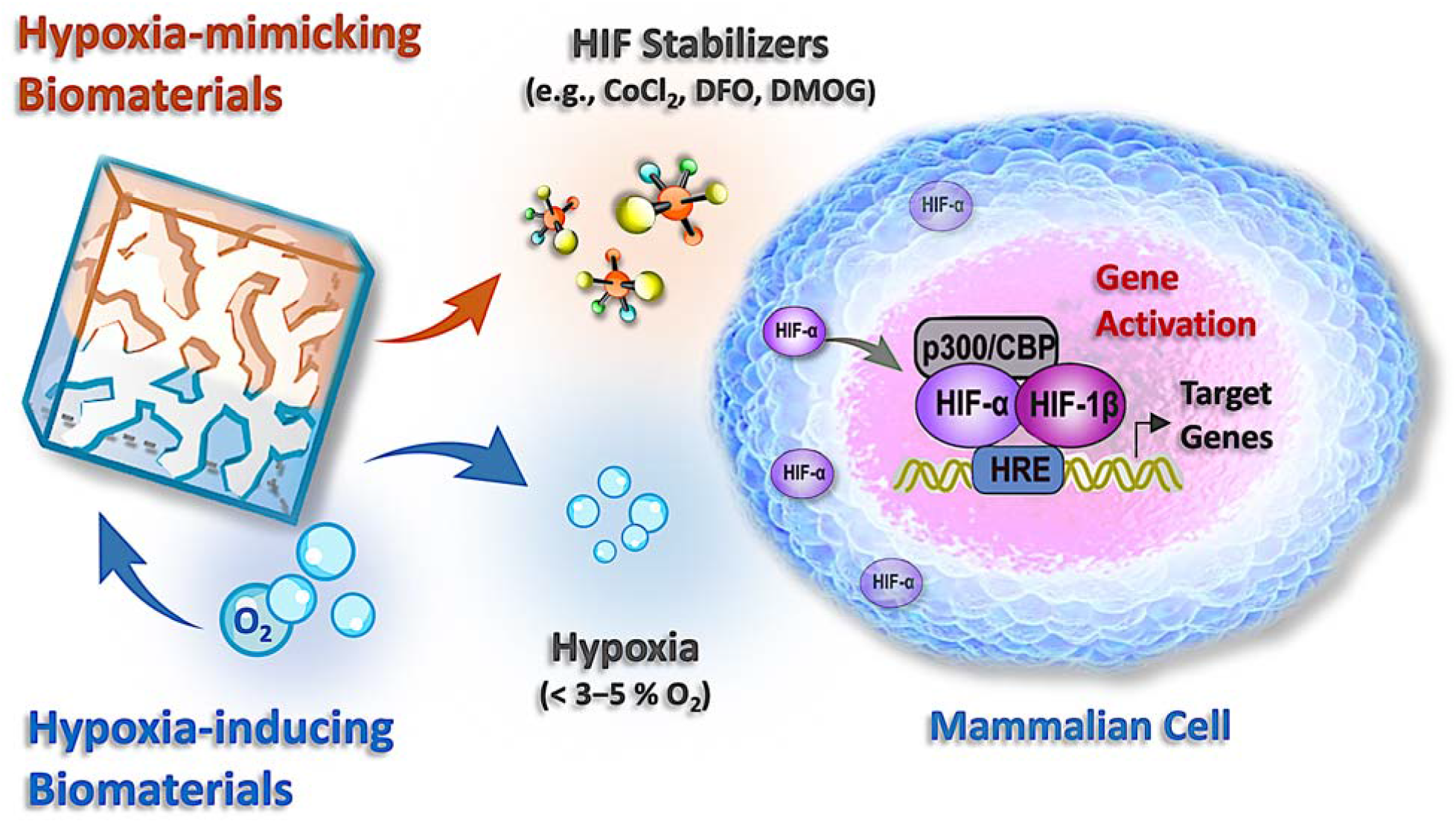

